# Endogenous Mechanical Reporters Encode the Global Mechanical Landscape of Morphogenesis

**DOI:** 10.64898/2026.11.08.765212

**Authors:** Ruoqing Xu, Han Gao, Fancheng Li, Lei-han Tang, Hai Zhu, Zhennan Zhou, Hanqing Guo

**Affiliations:** Center for Interdisciplinary Studies, Westlake University, Hangzhou, Zhejiang, China; Institute for Theoretical Sciences, Westlake University, Hangzhou, Zhejiang, China

## Abstract

Developmental systems usually comprise multiple genetically programmed morphogenetic events within a mechanically continuous environment, yet how individual processes interact and contribute to the overall mechanical landscape remains unknown. Here, we introduce the concept of “mechanical reporter”: endogenous tissues function as *in situ* probes of the embedded mechanical landscape. We exemplified cephalic furrow (CF) migration as a mechanical reporter in gastrulating *Drosophila*. Using optogenetics and genetics, we revealed that CF displacement reflects the combined effect of both force and tissue mechanical properties. By incorporating condition-specific mechanical landscape into a continuum minimal-response model, we developed a framework to quantitatively predict the reporter dynamics. Using different genetic mutants, we successfully decoded changes in the mechanical landscape from CF dynamics in gastrulating *Drosophila*. Overall, we established a predictive framework that uses CF dynamics to infer changes in the global mechanical landscape. Beyond this, the concept of mechanical reporter provides a general strategy for systematically understanding and decoding complex mechanical environments in living systems.

## Introduction

A fundamental challenge in biology is understanding how multiple genetically programmed modules in a mechanically continuous environment contribute to the global behavior. Patterned gene expression can generate tissue-intrinsic forces that drive individual morphogenetic events, such as folding and elongation^1,2^. This can be viewed as imposing a locally preferred mechanical landscape, such as a target curvature or strain, and pushing the tissue toward it. Yet in complex scenarios like embryogenesis, these processes occur simultaneously within a shared mechanical environment, such that forces generated in one tissue couple to and reshape others. Accordingly, the global morphology is not simply the sum of independent morphogenetic programs, but emerges from inter-tissue mechanical integration: gut looping arises from differential growth between the epithelium and the surrounding mesentery^3^ and zebrafish epiboly requires the yolk syncytial layer to pull the enveloping layer^4^. Despite growing recognition of inter-tissue mechanical coupling, quantitative decoding of this integrated mechanical landscape *in vivo* remains underdeveloped.

Recent studies largely address this problem in a forward direction. Genetic or mechanical perturbations target a specific morphogenetic process and illustrate how individual processes influence the mechanical landscape, as revealed by methods such as particle image velocimetry and strain mapping^5,6^. These approaches are powerful in establishing causal contributions, but the complementary reverse problem remains underexplored: whether the integrated mechanical landscape can be extracted from the dynamics of the endogenous tissue.

We propose that this problem can be resolved by a common feature of morphogenetic systems: tissues often continue to respond to external stresses after the completion of their genetically programmed morphogenesis^7–11^. These post-morphogenetic dynamics therefore encode information about the global mechanical landscape. Systematically monitoring the post-morphogenetic dynamics of such tissue, which functions as an endogenous **mechanical reporter**, could link these dynamics to individual force components and reveal mechanical factors that are hard to measure directly, such as mechanical properties, much like molecular reporters reveal hidden regulatory interactions. To qualify as a mechanical reporter rather than merely a responsive structure, a tissue should respond sensitively to the changing mechanical landscape and, conversely, the landscape should be inferable from the dynamics of the reporter, such that its behavior can be predicted under independent perturbations. This general framework should, in principle, apply to any post-morphogenetic tissue embedded in a mechanically continuous environment.

The gastrulating *Drosophila* embryo offers a unique opportunity to provide proof of principle for this concept. Multiple morphogenetic programs, including cephalic furrow (CF) folding^12–14^, ventral furrow (VF) internalization^15–18^, germ band extension (GBE)^19–22^, and posterior midgut (PMG) invagination^23,24^, proceed simultaneously within a shared mechanical environment. Therefore, these morphogenetic events are mechanically coupled: VF folding is facilitated by surrounding GB^25,26^, VF folding slows AP cell elongation during GBE^27^, and GBE is promoted by PMG invagination^28–30^. Among these events, we identified the CF as a candidate structure containing processes that function as mechanical reporters. Positioned at the head-trunk interface, it occupies a geometrically central location where forces from multiple morphogenetic processes converge^31,32^. Following its active actomyosin-driven folding phase, we showed that the CF undergoes substantial anterior-posterior displacements while large-scale morphogenetic events occur around it, consistent with previous observations^33^. The dorsal and ventral regions of the same furrow move in opposite directions, providing two spatially separated yet mechanically coupled response channels within a single structure.

By systematically perturbing the surrounding tissues and quantifying post-folding CF dynamics, we found that the dorsal and ventral CF channels read distinct yet coupled aspects of the global mechanical landscape. In particular, the dorsal CF is sensitive to the propagated expansion force generated by head cell mitosis, and the ventral CF is sensitive to the large tissue deformations like folding and elongation, both of which are influenced by the tissue mechanical property. These CF dynamics form a quantitative relationship with changes in the global mechanical landscape. We used mechanical landscape changes, such as differences in force-generating processes or mechanical properties, as the input, combined with geometrical confinements to construct a continuum minimal-response model, and predicted the dorsal and ventral CF trajectories via global energy minimization. After calibrating the relation between mechanical landscape changes and CF dynamics, conversely, we successfully identified changes in the global mechanical landscape using CF displacements across different genetic mutants.

Together, our results establish CF migration as an endogenous mechanical reporter and provide a strategy for future inter-tissue mechanics studies in the gastrulating *Drosophila* embryo: changes in the integrated mechanical environment are encoded in post-folding CF dynamics. More broadly, this study provided an example to show that the global mechanical landscape can be read through the dynamics of the tissue embedded within it. Any tissue that has completed its active intrinsic morphogenesis but remains mechanically coupled to its surroundings might serve as an endogenous probe. In this way, the *in situ* mechanical reporter concept offers a route towards a systems-level understanding of mechanical integration in complex environments.

## Results

### External-force-dominated post-folding CF migration qualifies as a mechanical reporter

The CF is traditionally recognized as a prominent epithelial fold that forms during *Drosophila* gastrulation and separates the anterior head from the trunk^31,32^. Through long-term 3D live imaging, we reconstructed the temporal progression of CF dynamics, revealing two distinct stages: an initial CF propagation period, during which the furrow folds and extends along the dorsal-ventral axis, followed by a CF migration period (Fig. 1a, d). During its initial propagation stage, actomyosin contractility is exerted along the lateral side of the CF cells to achieve an out-of-the-plane invagination towards the interior of the embryo^14^ (Fig. 1a). This active folding establishes the mature CF structure and defines its initial geometry. After completion of this active folding process, the CF undergoes an extended anterior-posterior displacement that is perpendicular to the direction of intrinsic actomyosin contractility^33^ (Fig. 1a). This post-folding movement, therefore, cannot be driven by the intrinsic contractile machinery of CF. The folded CF exhibited an anisotropic folding pattern during the migration phase, suggesting that it is being shaped by different external stresses (Fig. 1a, b). Consistently, recent findings showed that in mutants lacking CF structure, global compressive stresses result in ectopic folds at the head-trunk boundary, at a position analogous to that of the wild-type CF^12,13^. Together, these observations indicate that the post-folding displacement of the CF is governed predominantly by external forces and that CF migration might serve as an endogenous mechanical reporter of gastrulation in the *Drosophila* embryo (Fig. 1d).

**Fig. 1.**
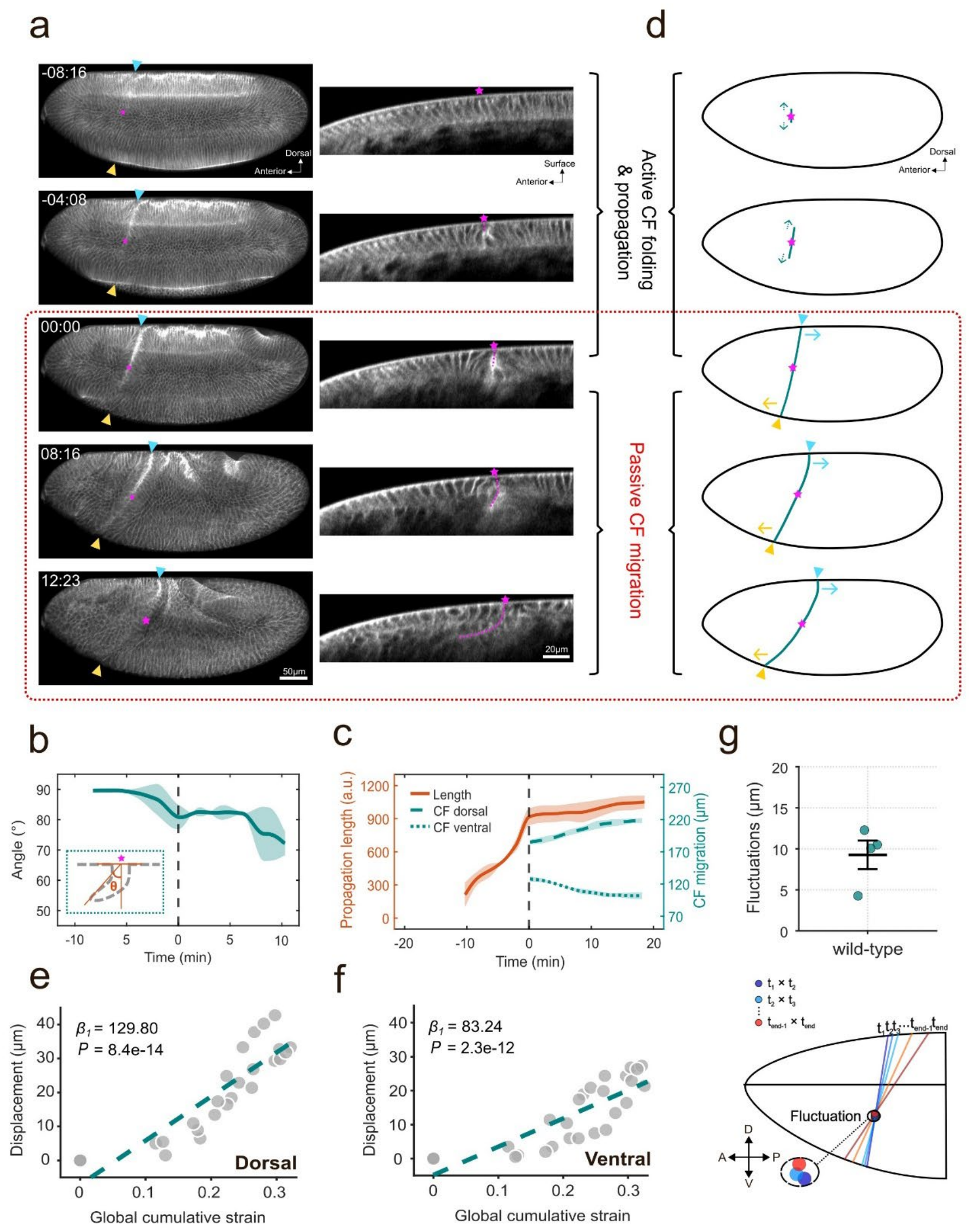
The cephalic furrow migration as a passive mechanical reporter. **(a)** CF dynamics in wild-type *Drosophila* embryos during gastrulation. CF folding initiated at the lateral side of the embryo (magenta star) and propagated towards the dorsal and ventral sides (left, cyan and yellow arrowheads, respectively). 0 min marks the onset of CF migration in the following figures, unless noted differently. Folded CF was labeled by a dotted magenta line in the lateral view (right). **(b)** Angle of CF folds over time. The angle reports the anisotropy of the folded CF during the migration stage, indicating that the furrow was shaped by differential external stresses. Inset: angle definition. Mean ± s.d.; n = 2 embryos. **(c)** CF propagation and migration distance over time. n = 2 embryos. **(d)** Schematic of the two-stage CF dynamics. **(e, f)** Instantaneous dorsal **(e)** and ventral **(f)** CF displacement plotted against the embryo’s global cumulative strain (relative to T_0_). Dashed line: linear regression (linear mixed-effects model); Dorsal: *P* = 8.4×10^-14^, *β*_*1*_ = 129.80, 95% CI: [109.71, 149.89]; Ventral: *P* = 2.3×10^-12^, *β*_*1*_ = 83.24, 95% CI: [68.49, 97.99]; n = 3 embryos. **(g)** Fluctuations of the rotational center. The schematic illustrates the measurement design. See also Fig. S1. Mean ± s.d.; n = 4 embryos.

To systematically monitor the CF response to external stresses, we tracked the most dorsal and most ventral regions of the folded CF (dorsal CF and ventral CF, respectively) as indicators of post-folding displacement along the anterior-posterior axis. We defined T_0_ as the time at which

CF propagation ends, and the dorsal and ventral CF tips become visually distinguishable (Fig. 1c). In wild-type embryos, we found that the dorsal CF moves progressively towards the posterior, whereas the ventral CF moves towards the anterior (Fig. 1a, c). The bidirectional CF movements exhibited strong positive correlations with the embryo’s global cumulative strain (Fig. 1e, f), indicating that CF displacements quantitatively reflect changes in the global mechanical landscape. Notably, the trajectories of dorsal and ventral CF movements converged on a defined, coherent rotational center that was maintained over time (Fig. 1g, Fig. S1), suggesting that dorsal and ventral CF dynamics are mechanically intertwined and coordinated at the organism level.

### Dorsal and ventral CF channels report distinct mechanical sources

To evaluate whether CF migration could serve as a mechanical reporter, we first perturbed the major force-generating events during the post-folding phase to assess its sensitivity to changes in the mechanical landscape. During this period, the principal force-generating mechanisms are mitosis-related expansion and actomyosin-driven morphogenetic events, including GBE and PMG invagination^22,24,34^ (Fig. 2a). Although the initiating cells of the anterior and posterior dorsal folds (ADF and PDF) undergo a local increase in deformability^35,36^, the folds are established by surrounding tissue^37^. The anterior midgut (AMG) is driven by single-cell level changes that have limited contribution to large-scale tissue displacements^38^ (Fig. 2a). Thus, we focused on mitotic expansion and GBE-PMG as the major candidates responsible for generating embryo-scale mechanical inputs during CF migration.

**Fig. 2.**
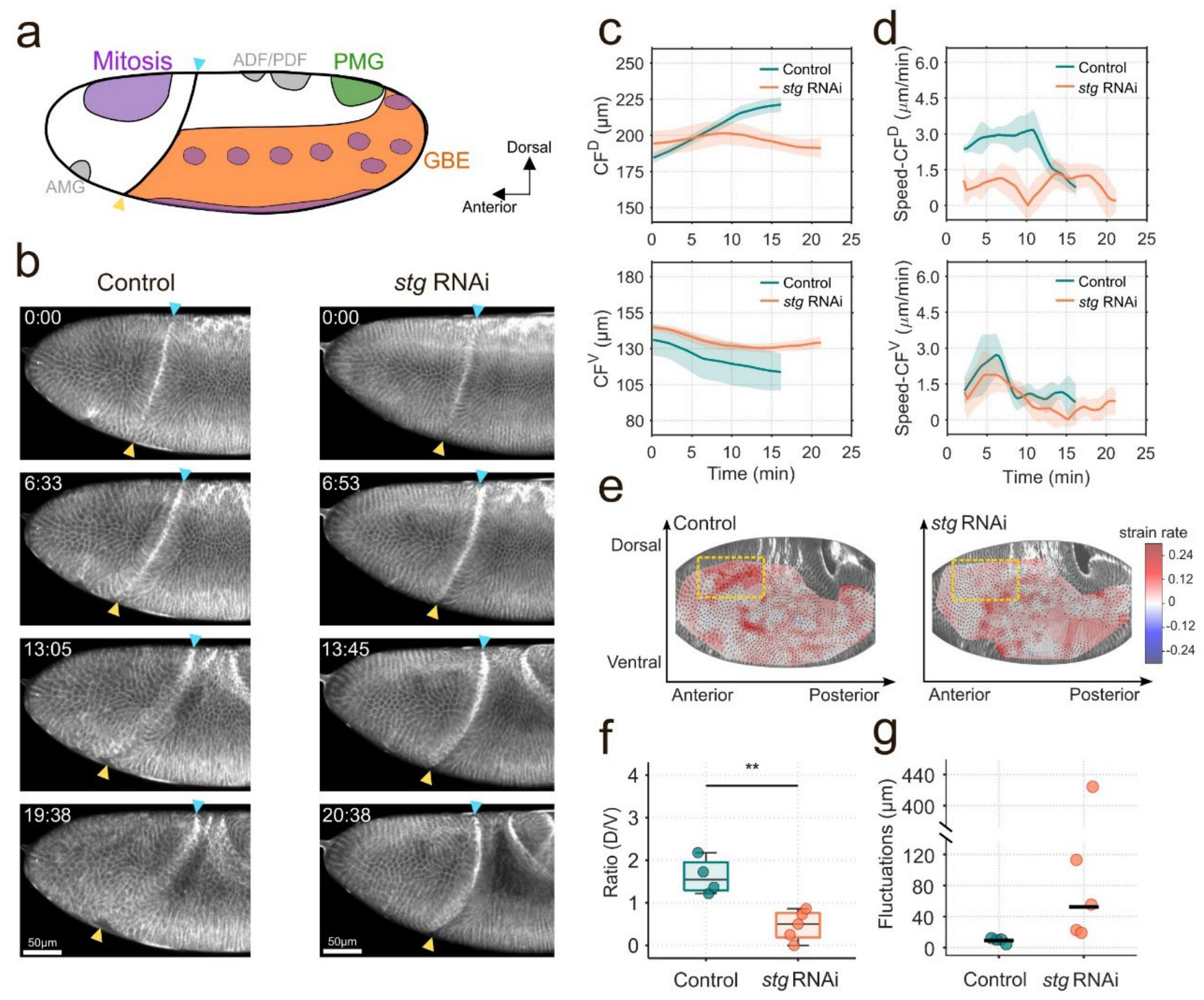
Dorsal CF displacement reflects the head mitosis activity. **(a)** Schematic of the morphogenetic events during CF migration. Colored regions (purple, green, and orange) decode the major force contributors in gastrulating *Drosophila*. **(b)** Representative *en face* views of wild-type and *stg* RNAi embryos during CF migration. Cyan and yellow arrowheads mark the dorsal and ventral CF, respectively. **(c, d)** Dorsal (top) and ventral (bottom) CF displacements **(c)** and velocities **(d)** over time in wild-type and *stg* RNAi embryos. **(e)** Strain maps of wild-type and *stg* RNAi embryos during CF migration. The dashed yellow box outlines the major mechanical difference in the dorsal head region, consistent with the loss of mitotic expansion in the head. **(f)** Ratio of dorsal-to-ventral CF displacement in wild-type and *stg* RNAi embryos. *t*(7) = 4.47, *P* = 2.9×10^-3^; Cohen’s *d* = 3.00, 95% CI: [2.36, 6.42]. **(g)** Rotational center fluctuations in wild-type and *stg* RNAi embryos. Horizontal lines denote the mean values. Mean ± s.d.; n = 4 (wild-type) and 5 (*stg* RNAi) embryos.

First, to assess the contribution of mitotic activity, we used *string* (*stg*) RNAi to inhibit cell division^39,40^. Compared with wild-type embryos, *stg* RNAi embryos exhibited markedly slower dorsal CF movement, whereas ventral CF movement remained relatively comparable, with only a minor reduction (Fig. 2b, c, d). In wild-type embryos, mitosis occurs in a genetically programmed spatiotemporal pattern, with several head mitotic domains positioned anterior to the dorsal CF (Fig. 2a). Strain mapping further revealed that the major mechanical difference between *stg* RNAi and wild-type embryos occurs in the head region (Fig. 2e), where mitosis-associated tissue deformation was strongly reduced, indicating a specific loss of tissue expansion generated by head mitosis. Importantly, mitotic domains in the trunk during the CF migration window did not appear to influence CF movement (Fig. 2a, c, d). Due to the strongly reduced dorsal CF migration in *stg* RNAi embryos, the displacement ratio between dorsal and ventral CF is reduced (Fig. 2f), and the rotational center is shifted close to the dorsal side and becomes dominated by noise (Fig. 2g), indicating that the coherent rotational geometry depends on the balance between dorsal and ventral mechanical factors.

Next, to evaluate how actomyosin-driven morphogenesis contributes to the global mechanical landscape, we used an optogenetic tool to inhibit actomyosin contractility in GBE and PMG^26^ (Fig. 3a). Notably, mitosis still occurs in the optogenetically-treated embryos (Fig. S2, white circle and box). Reducing GBE-PMG activity resulted in a pronounced decrease in ventral CF velocity, suggesting that GBE-PMG force is important for ventral CF displacement (Fig. 3b, d, f, OptoL). Yet it is not the only source of forces, as the ventral CF continues to migrate (Fig. 3f). To further explore the nature of the other contributors, we perturbed actomyosin activity in embryos at the end of cellularization, before VF folding, GBE, and PMG occur (Fig. 3a-c, OptoE). Without any large tissue deformation, the ventral CF showed minimal displacement (Fig. 3e). These observations together suggest that VF influences CF migration dynamics, despite the fact that VF folding is completed before the onset of CF migration. Tissue flow analysis of embryos during and immediately after VF folding revealed a comparable vortex pattern (Fig. 3g, red box). Furthermore, strain mapping of embryos during VF folding revealed higher strain in the surrounding GB, consistent with previous studies^41^. Notably, this pattern, as well as the ventral-directed tissue flow, persists in embryos that have completed VF folding (Fig. 3g and 3h, cyan box). The data revealed that large tissue deformations, such as VF, establish sustained tissue flow that outlasts the morphogenetic event that created them, thereby contributing to the global mechanical landscape. Overall, we showed that dorsal CF is sensitive to head mitosis, and ventral CF is sensitive to GBE-PMG forces and VF flow.

**Fig. 3.**
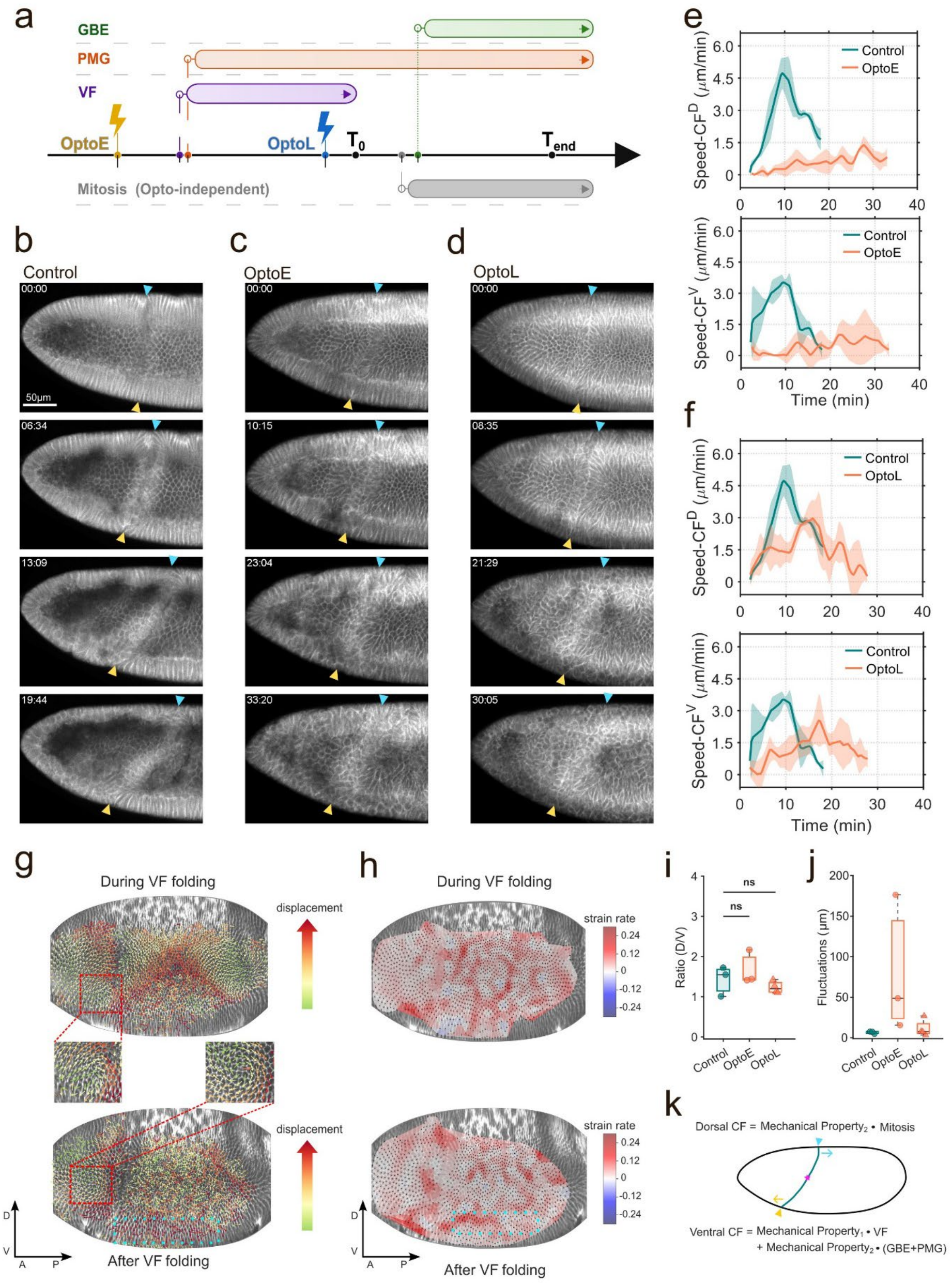
CF migration reflects both morphogenetic forces and the mechanical property. **(a)** Developmental timeline of the morphogenetic events before and during CF migration and the experimental perturbation strategy. OptoL group inhibited GBE-PMG with VF folding intact, whereas the OptoE group inhibited both VF and GBE-PMG. **(b-d)** Representative *en face* views of wild-type **(b)**, OptoE **(c)**, and OptoL **(d)** embryos during CF migration. **(e-f)** Dorsal and ventral CF velocities over time in wild-type and OptoE embryos **(e)** and OptoL embryos **(f). (g-h)** Tissue flow fields **(g)** and strain map **(h)** of wild-type embryos during and immediately after VF folding. The two windows exhibited similar flow patterns (see the red box). Elevated strain rate in the GB during VF folding persisted after folding was complete, together with sustained ventral-directed tissue flow (cyan box). Both observations suggested that VF folding established a tissue flow that outlasted the event itself. **(i)** Ratio of dorsal-to-ventral CF displacement in wild-type, OptoE and OptoL embryos. *F*(2, 7) = 1.64, *P* = 0.26 (One-way ANOVA). **(j)** Rotational center fluctuations in wild-type, OptoE and OptoL embryos. **(k)** Dorsal and ventral CF channels reported distinct yet coupled aspects of the global mechanical landscape. Mean ± s.d.; n = 3 (wild-type), 3 (OptoE), and 4 (OptoL) embryos.

### CF migration reflects both morphogenetic forces and the mechanical property

Interestingly, the optogenetic perturbation affects not only the ventral but also the dorsal CF in both groups (Fig. 3e, f). This observation was unexpected because: (1) as the major driving force for dorsal CF displacement, mitosis is not influenced; and (2) GBE-PMG-generated forces are oriented in the opposite direction to dorsal CF displacement. If the dorsal CF also responds to local pushing or pulling generated by GBE-PMG, its inhibition should accelerate the dorsal CF displacement. The fact that dorsal CF was slowed instead indicates that the perturbation altered a global mechanical factor beyond the local driving force itself.

The dorsal CF displacement can be interpreted as local tissue flow in the dorsal CF region. Conceptually, tissue flow follows the force-velocity constitutive law and is determined by the integrated effects of driving force and the mobility, which, in our case, can be approximated by the mechanical property of the tissue^42^. Therefore, both the dorsal and ventral CF displacements are influenced by the viscoelastic property of the tissue, despite the differences in the driving forces. The observation that dorsal CF was slowed in both OptoE and OptoL groups, even though head mitotic activity was not directly targeted (Fig. 3e, f), indicates that the tissue mechanical property might be influenced. The optogenetic tool inhibits both actomyosin contractility and the actin network^26^, which is a major contributor to tissue mechanical property^43^. It is therefore possible that the mechanical property is also influenced by optogenetic treatments. Consistent with this interpretation, the ratio between dorsal and ventral CF displacement in the OptoE, OptoL, and the wild-type embryos showed a statistically non-significant change, possibly because the alteration of mechanical property masked the influence of lacking large tissue deformations (Fig. 3i). More interestingly, the fluctuation in the rotational center becomes larger in the OptoL group, and further increases in the OptoE condition, possibly dominated by intrinsic noise, indicating that both driving forces and tissue mechanical property govern the global mechanical landscape stability (Fig. 3j).

Together, the dorsal CF displacement is the integration of mechanical property and mitosis, and the ventral CF displacement is affected by VF-induced tissue flow, GBE-PMG force, and tissue property (Fig. 3k). The mechanical property thereby couples the dorsal and ventral CF displacements. Collectively, the data support CF migration as the mechanical reporter for gastrulating *Drosophila*, and reveal that post-morphogenetic ventral and dorsal CF displacements encode distinct yet coordinated components of the global mechanical landscape.

### A continuum minimal-response model maps mechanical inputs onto CF dynamics

After experimentally establishing the CF migration as a mechanical reporter, we seek to quantitatively understand the input-output relationship between the global mechanical landscape and CF dynamics. We used a continuous elastic curve, constrained by the eggshell *U*_conf_, to mimic the mid-sagittal cross-section of the embryo (Fig. 4a, blue curve). The anterior pole in the model is stabilized by applying a soft-point penalty *U*_pin_ (Fig. 4a, green square), as evidenced by experimental observations (Fig. S3a). A preferred-curvature field, *κ*_0_(*s, t*), is adapted in the model to mimic the genetically patterned CF position and furrow morphology during the reporter phase. The changes in the local strain fields are reconstructed by building a condition-specific target stretch field *g*(*s, t*), which is composed of stretching zones based on the biological processes (mitosis, GBE-PMG, and VF flow) that serve as the mechanical contributors to the reporter dynamics (Fig. 4a). At each time point, the profile of *g*(*s, t*) is updated according to the presence of different biological events. Thus, the dynamic tissue configuration is determined under a quasi-static approximation by minimizing the total mechanical energy (Eq. (1)):

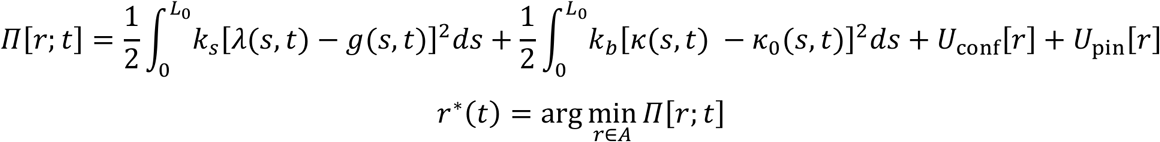

**Fig. 4.**
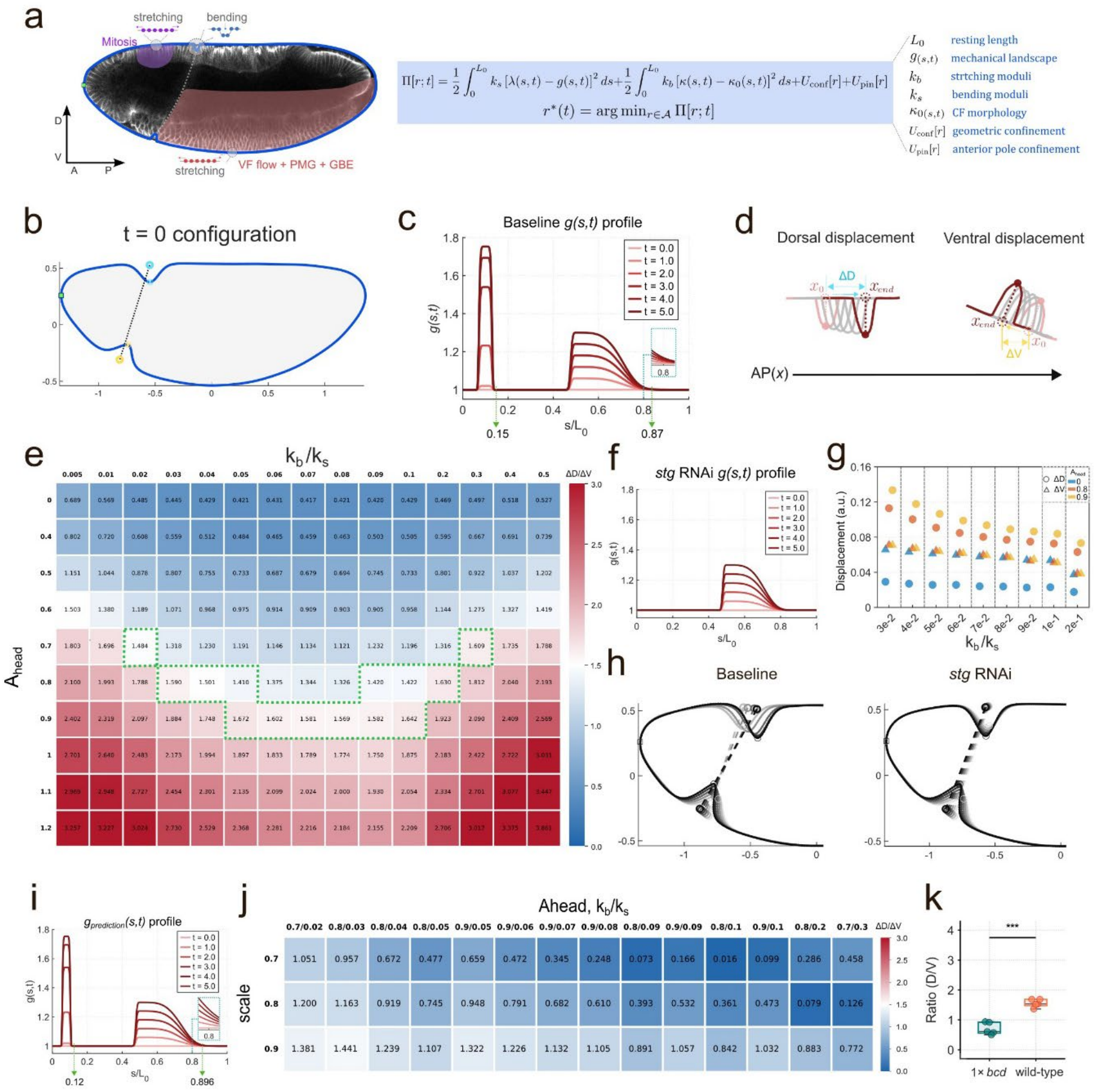
A continuum minimal-response model recapitulates mechanical reporter dynamics. **(a)** Model design. The dynamic embryo configuration is determined by stretching and bending energies, geometric confinement, and a soft-point penalty. **(b)** The emerged t = 0 configuration when g(s,0) = 1. Parameters were determined by matching this shape to the wild-type embryo morphology provided in (a). **(c)** The global mechanical landscape *g*_*baseline*(*s,t*)_ over time for wild-type embryos. Green arrows mark the CF position. **(d)** The calculation of dorsal and ventral CF displacement. **(e)** The heatmap of 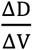 under different combinations of *A* _*head*_ and *k*_*b*_/ *k*_*b*_. **(f)** The example mechanical landscape profile for *stg* RNAi embryos at *A*_*head*_ = 0.8 and *k*_*b*_/*k*_*s*_ = 0.04. **(g-h)** The model recapitulated the reporter’s dynamics in the *stg* RNAi condition. Example morphology shown here is *A*_*Prediction*(*s,t*)_ = 0.8 and *k*_*b*_/*k*_*s*_ = 0.04. **(i)** The global mechanical landscape *g*_*Prediction*_ over time. **(j)** The heatmap of 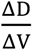 across different proportionally anterior shifts of CF under proper combinations from **(e). (k)** Ratio of dorsal-to-ventral CF displacement in wild-type and 1x *bcd* embryos. *t*(8) = -8.06, *P* = 4.1×10^-5^; Cohen’s *d* = -5.10, 95% CI [-15.35, -4.15]. n = 5 (wild-type), 5 (1x *bcd*) embryos.

Here, *λ*(*s, t*) denotes the local stretch ratio; *k*_*s*_ and *k*_*b*_ reflect the tissue tendency to stretch or bend, respectively; *κ*(*s, t*) denotes the local curvature of the tissue curve and reflects the current CF position and morphology. The admissible configuration space A contains closed curves that remain within the confining geometry. The detailed model formulation, parameterization, and numerical implementation are provided in the Materials and Methods section. Parameters were determined by recapitulating the wild-type embryo morphology at the onset of CF migration (defined as *t* = 0, *g*(*s*, 0) = 1, Fig. 4b). During CF migration, the mechanical landscape, *g*_*basline*_ (*s, t*), changes over time and is determined by the morphogenetic events in a wild-type embryo (Fig. 4c). The model-emerged dorsal and ventral CF displacements were then calculated, and the ratio 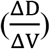 was quantified (Fig. 4d).

The CF displacement in the model is sensitive to (1) the spatial balance of the force, represented by the ratio between head stretching zone, *A*_*head*_, and the body stretching zone; and (2) the mechanical property preference to bending or stretching, represented by the ratio between *k*_*b*_ and *k*_*s*_ . Through calibrating the simulated 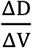 with the wild-type embryo dynamics (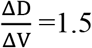, Fig. 2f), we defined the proper combinations of *A*_*head*_ and 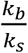 for the baseline condition (Fig. 4e, green box). Then, to mimic the mechanical effect of *stg* RNAi, we modified the *g*_*mitosis*_ (*s, t*) by reducing *A*_*head*_ to 0 (Fig. 4f). The model successfully recapitulated the CF dynamics observed in the *stg* RNAi condition, in which dorsal CF no longer migrates over time (Fig. 4g, h).

To evaluate the predictability of this continuum minimal-response model on capturing the dynamics of mechanical reporters, we altered the position of CF. Specifically, the prescribed dorsal and ventral fold centers in *κ*_0_(*s, t*) were shifted anteriorly, and *g*_*prediction*_ (*s, t*) was modified accordingly (Fig. 4i). Other parameters were held fixed at their baseline values. Simulated results showed that the anterior shifting of CF resulted in a reduction of the 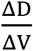 across different *A*_*head*_ and 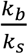 combinations (Fig. 4j). To experimentally validate this model prediction, we quantified the CF dynamics in embryos from *bicoid* heterozygous mutant mothers (1x *bcd*), in which the CF forms at a more anterior position^31^ (Fig. S3b). Consistent with the model prediction, the 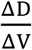 in 1x *bcd* is markedly reduced (Fig. 4k). Therefore, we successfully established a continuum minimal-response model that maps the changes in mechanical landscape onto the CF dynamics, further supporting the idea that CF migration functions as an *in situ* mechanical reporter for gastrulating *Drosophila*.

### Decoding mechanical landscape changes using CF dynamics

Having established that CF migration encodes the global mechanical landscape, we next asked if the relationship could be inverted: whether an unknown change in the mechanical landscape could be decoded from the reporter dynamics. As illustrated previously, the CF dynamics in the 1x *bcd* embryo differ from those in wild-type embryos, but the change in the global mechanical landscape remains unknown. The ventral CF in 1x *bcd* is unaffected, indicating the preservation of the ventral CF-relevant response between wild-type and 1x *bcd* embryos. Since the large tissue deformations, such as VF/GBE/PMG, are not influenced by the treatment, the data suggest that the mechanical property of the tissue is likely comparable between wild-type and 1x *bcd* embryos (Fig. 5a). Then the reduction in the dorsal CF displacement in 1x *bcd* embryos is likely due to changes in head mitosis (Fig. 5b). Because mitotic patterning is unaffected in 1x *bcd* embryos, the reduced head expansion most likely reflects a smaller pool of mitotic cells rather than a reduced mitosis activity. Using 3D segmentation, we confirmed that 1x *bcd* embryos contain fewer head cells (Fig. 5c), suggesting that changes in the mechanical landscape could be decoded from CF dynamics.

**Fig. 5.**
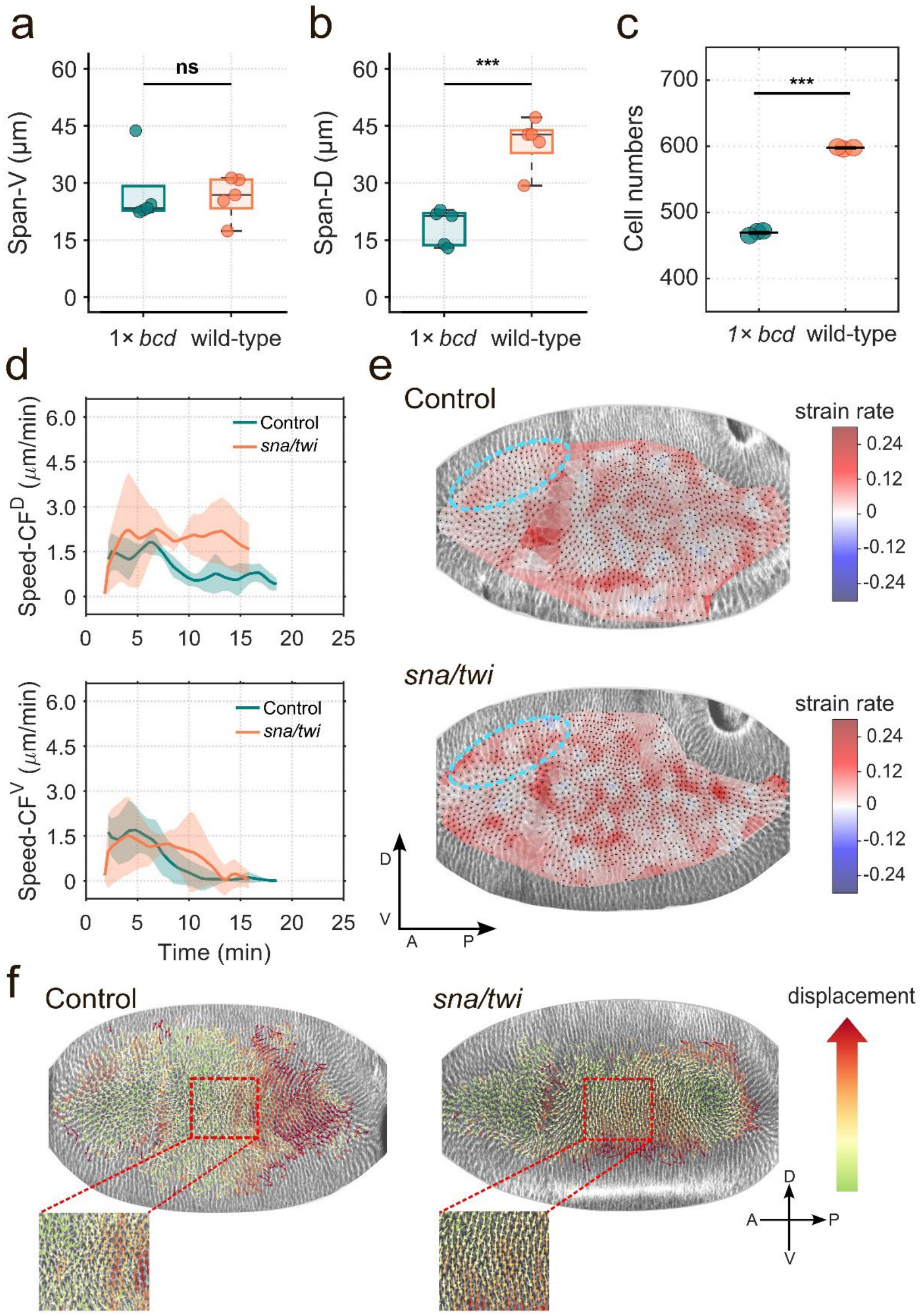
CF dynamics decode the mechanical landscape in gastrulating *Drosophila*. **(a-b)** Ventral **(a)** and dorsal **(b)** CF displacements in wild-type and 1x *bcd* embryos. n = 5 (wild-type), 5 (1x *bcd*) embryos. *t*(8) = 0.09, *P* = 0.93 (ventral CF displacements); *t*(8) = -5.84, *P* = 3.88×10^-4^, Cohen’s *d* = -3.69, 95% CI [-10.25, -2.45] (dorsal CF displacements). **(c)** Head cell numbers in wild-type and 1x *bcd* embryos. n = 3 embryos. *t*(4) = -54.45, *P* = 6.8×10^-7^; Cohen’s *d* = -44.46, 95% CI [-312.72, -41.71]. **(d)** Dorsal (left) and ventral (right) CF displacement velocities over time in wild-type and *snail/twist* mutant embryos. **(e)** Strain maps of wild-type and *snail/twist* embryos at T_0_. VF loss remodeled the mechanical landscape and resembled the control strain map, but with increased strain in the anterior-dorsal region as indicated by the cyan circle. **(f)** Tissue flow fields of wild-type and *snail/twist* embryos during VF folding. Color scale denotes displacement magnitude (identical limits for both genotypes). Ventral-directed flow is highlighted by the red box. Mean ± s.d.; n = 4 embryos per condition.

To further investigate whether the CF migration reports the integrated mechanical landscape rather than the identity of individual force-generating processes, we tracked CF dynamics in the *snail/twist* mutants^44^. These embryos lack the VF, a major morphogenesis event. Notably, dorsal CF displacement was accelerated in these embryos, whereas the ventral CF displacement was comparable to that of a wild-type embryo (Fig. 5d). These CF dynamics indicate that the global mechanical landscape should be comparable between these genotypes, with an enhanced mechanical input anterior to the dorsal CF. Strain mapping confirmed the reporter-based inference and showed that the head region of *snail/twist* mutants exhibited enhanced strain relative to a wild-type embryo (Fig. 5e, cyan circle). The overall global mechanical landscape at the onset of the reporter phase, particularly in the trunk region, is otherwise similar, as indicated by tissue flow and strain mapping (Fig. 5e, f). In this condition, the force source shifted from VF folding to global stress, as indicated by the reported GBE-like cell movements in VF mutants^26^, but the integrated global mechanical landscape remained comparable except for the enhanced head strain. Together with the 1x *bcd* analysis, these results establish that using post-folding CF dynamics, we could decode unknown changes of the global mechanical landscape in the gastrulating *Drosophila*.

## Discussion

In this study, we proposed that post-morphogenetic tissue dynamics could serve as a mechanical reporter of the global mechanical landscape. Using the gastrulating *Drosophila* embryo as an example, we established that CF migration is governed predominantly by external forces and can serve as an *in situ* mechanical reporter. Through experimental perturbations, we showed that CF displacements reflect the combined effects of mechanical properties and tissue forces. Specifically, the dorsal CF is sensitive to the head mitosis, and the ventral CF is more sensitive to the actomyosin-driven tissue deformations. More interestingly, our experimental inhibition of actomyosin contractility revealed a hidden component: tissue flow induced by VF folding persists beyond the folding event itself, shaping the global mechanical landscape during CF migration. The continuum minimal-response model we established here showed that CF displacement emerges once compatibility between tissue-intrinsic strain fields and global geometric constraints is achieved. Consequently, an identical local perturbation can produce different CF responses when the surrounding mechanical landscape, tissue geometry, or confinement state differs. By treating the post-folding CF as a mechanically responsive probe embedded in this system, the model converts changes in the condition-specific mechanical landscape into quantitative predictions of CF trajectories. In general, this formulation provides a framework for connecting local tissue remodeling with global morphogenetic outcomes in confined epithelial systems. In systems dominated by constriction or pulling rather than compression and stretching, the same framework can be extended by incorporating the corresponding mechanical inputs.

The mechanical reporter strategy occupies a unique methodological niche among existing approaches. Model-based force inference methods extract forces from cell shapes and motions, but rely on assumed constitutive relations^45^. The mechanical reporter calibrates the relationship empirically through biological perturbations, so the effective response relation is measured in the embryos’ native state. Engineered probes, such as oil droplets^46^ and ferrofluid droplets^47^, yield absolute tissue-level stresses but are often limited by their invasiveness and by the fact that they report only the mechanical state in the immediate vicinity of the probe. The mechanical reporter uses endogenous tissue as the probe and reports the integrated effect of the propagating forces, at the cost of relative rather than absolute measurements. Morphogenetic behaviors have long served as qualitative stress readouts, from the wrinkling instability in brain development^48^ to the ectopic folds that arise when the CF is absent^12,13^. The present framework extends this logic from static, qualitative readouts to a quantitative, dynamical readout from an endogenous tissue in its native state.

Beyond the gastrulating *Drosophila* embryo, the mechanical reporter framework should be applicable to many other biological systems. A tissue would qualify as a mechanical reporter if: (1) its genetically programmed active morphogenesis is largely complete before the reporter phase begins, or the active contribution can be calibrated or is negligible, such that the reporter dynamics that are driven by extrinsic stresses can be quantified; (2) it is experimentally accessible for establishing the reproducible, calibratable reporter responses under different global mechanical landscapes; (3) the relation between changes in the mechanical landscape and the reporter responses should be invertible, so that the changes of the mechanical landscape can be inferred from the reporter dynamics. For instance, in zebrafish, the anterior neural plate exhibits low intrinsic force and is reshaped by global tissue flows at the late gastrula stage^49^, making it a potential mechanical reporter of the zebrafish gastrulation after proper calibration. Furthermore, the nature of a mechanical reporter is not necessarily limited to the tissue; any passive interface may serve the purpose. For instance, common tissue-folding processes, such as neural tube closure in vertebrates or dorsal closure in *Drosophila*, result in a seam on the embryonic surface that is subject to external loads^50,51^. The shape of this passive interface could reflect the surrounding compression and serve as a mechanical reporter for the orthogonal mechanical landscape. This framework may also benefit organoid studies when 3D morphogenetic events such as budding and branching emerge stochastically, producing substantial sample-to-sample heterogeneity^52,53^. An *in situ* mechanical reporter would enable monitoring of the internal force balance and mapping of the global mechanical landscape in such systems, revealing hidden mechanical interactions and providing quantitative benchmarks needed to improve the reproducibility of organoid culture, especially under high-throughput conditions^54,55^.

The mechanical reporter framework also addresses a central problem in the physics of living systems. Tissues consume chemical energy to generate forces and deform far from thermal equilibrium, so any observed tissue motion integrates both passive mechanical responses and active, tissue-intrinsic forces^56^. Disentangling these contributions is the core difficulty for force inference in living systems^42^. Passive microrheology infers mechanical properties from the spontaneous motion of embedded tracers^57^, but converting these fluctuations into a quantitative response function requires the fluctuation-dissipation theorem, which fails when fluctuations are driven by molecular motors rather than thermal noise^58^. Active microrheology bypasses this problem by applying a known external force and measuring the response directly, but it perturbs the system being studied^59,60^. The mechanical reporter framework resolves this through experimental design rather than theoretical simplification: by selecting tissues with calibratable or negligible intrinsic activity, the measured reporter dynamics are dominated by extrinsic mechanical inputs. The relationship between reporter dynamics and the evolving global mechanical landscape can therefore be established via biological perturbations, conceptually analogous to measuring a system’s response function. By bridging the gene-centric, module-oriented view of developmental biology with the systems-level, continuum-oriented perspective of physics, the mechanical reporter’s framework offers a path toward a comprehensive understanding of inter-tissue mechanics that complements its molecular description.

## Materials and Methods

### Fly stocks and genetics

The following fluorescently marked fly lines were used in this study: Ecad-GFP^61^, Sqh-mCherry^16^, His2Av-RFP, 67 Sqh-GFP, Spider-*klar*-GFP, 15 P4M-mCherry, and 15 Gap43-mCherry. Opto-Rho1DN^26^ was used for optogenetic treatment. To inhibit mitosis, UAS-*stg* RNAi line (BDSC 36094, y[1] sc[*] v[1] sev[21]; P{y[+t7.7] v[+t1.8]=TRiP.GL00513}attP40) was crossed to 67 Sqh-mCherry;15 Ecad-GFP to generate UAS-*stg* RNAi/67 Sqh-mCherry;15 Ecad-GFP/+ females. To generate embryos lacking a ventral furrow, *snail*/*twist* double mutant (BDSC 3299, cn^1^ sna^18^ twi^3^ bw^1^ speck^1^/CyO) was used. To obtain embryos with a more anteriorly positioned CF, *bcd* mutants (Kyoto:107042, bcd^12^ p^p^/TM3, Sb^1^) were used.

### Live Imaging

Live imaging of *Drosophila* embryos was performed using an Olympus FV4000 two-photon microscope equipped with a 405/488-nm single-photon stimulation module. A 25× water-immersion objective (XLPLN25×WMP2, NA 1.05, WD=2 mm) was used for all experiments. Unless otherwise indicated, embryos were imaged after completion of cellularization using 920 nm and 1100 nm lasers. Imaging was performed in XYZT acquisition mode. Z-stacks were acquired from the embryo surface to a depth of approximately 100 μm, encompassing approximately one-half of the embryonic depth, with a z-step of 2 μm. The field of view was adjusted according to the region of interest, with an xy resolution of 0.497 μm/pixel. Successive Z-stacks were acquired at approximately 2 min intervals.

### Optogenetic Stimulation

Optogenetic perturbations were performed using the Opto-Rho1DN^26^ to acutely inhibit actomyosin contractility throughout the embryo at defined developmental stages. All embryo collection, handling, and sample preparation for optogenetic experiments were carried out in the dark under dim red illumination to avoid unintended activation of the CRY2/CIBN system.

During optogenetic experiments, time-lapse imaging was restricted to the 1100 nm laser to avoid unnecessary activation. Embryos were imaged before stimulation to establish a pre-stimulation baseline, then whole-embryo illumination was performed with 488-nm light at 5% laser power for 15 s. Two-photon time-lapse imaging was resumed thereafter to monitor tissue responses and CF dynamics. Successful optogenetic perturbation was assessed by changes in tissue morphology and dynamics.

### Quantification of CF migration dynamics

To quantify the spatiotemporal dynamics of post-folding CF migration, two reference time points were defined for each embryo. T_0_ was defined as the first frame in which the CF had extended to both the dorsal and ventral margins of the embryo, marking the completion of furrow propagation and the beginning of the migration phase. T_end_ was defined as the time point at which CF displacement reached its maximum and subsequently approached a short-term plateau.

Time-lapse images were analyzed in ImageJ (version 1.54p). For each embryo, the mid-sagittal plane containing the dorsal and ventral CF and the anterior pole was selected as the reference imaging plane. Images were preprocessed using Gaussian filtering (σ=1.5 pixels) followed by contrast-limited adaptive histogram equalization (CLAHE) to improve visualization. Using the ROI Manager, three landmarks were manually annotated in each frame: the dorsal CF, the ventral CF, and the anterior pole. Their two-dimensional coordinates were recorded and analyzed in MATLAB (R2024a) to generate time-resolved CF displacements and velocities.

### Quantification of CF rotational center stability

To quantify the fluctuation of the CF rotational center, we defined the instantaneous rotational center as the intersection between the lines connecting the dorsal and ventral CF at two consecutive time points. The coordinate of this intersection was tracked throughout the CF migration. Because CF displacement is subtle at early time points, the intersection of the two lines is geometrically ill-conditioned: a small measurement error in either CF position is amplified into a large error in the intersection coordinate, so these estimates are not reliable.

To obtain a stable, reproducible estimate, intersection coordinates from each embryo were partitioned using k-medoids clustering (*k* = 2, Euclidean distance, with fixed random seed). The main cluster was defined as the cluster containing ≥ 3 points. A cluster was classified as isolated and excluded only if all two of the following conditions were satisfied: (1) it contained ≤ 2 points, from the earliest accessible intersections ({T_0_-T_1_; T_1_-T_2_}); (2) its nearest-point distance to the main cluster exceeded the median within-main-cluster nearest-neighbor distance plus 2 × median absolute deviation (MAD).

Intersections remaining after this predefined filtering procedure were retained to quantify rotational-center fluctuations. Fluctuation was quantified as the range of the retained intersection coordinates along the AP axis. Smaller fluctuation values indicate greater spatial confinement and, consequently, higher stability of the CF rotational center during migration.

### Quantification of cell number in the dorsal head region

To quantify cell number in the dorsal head region, embryos expressing His2Av-RFP as a nuclear marker and Spider-*klar*-GFP as a membrane marker were imaged from the dorsal side using two-photon microscopy. Time-lapse Z-stacks covering approximately the dorsal half of the embryo were acquired from before gastrulation through early gastrulation. Cell number was quantified near the end of cellularization, thereby minimizing changes in tissue geometry. The position of the cephalic furrow is tracked and used to define the boundary of the head region. Regions of interest encompassing the dorsal head tissue were manually delineated across the corresponding optical sections, and the nuclei within the selected volume were segmented in three dimensions using Cellpose (v4.0.7)^62^ in Z-stack mode. Cell number was determined as the number of uniquely labeled nuclear objects within the segmented volume, excluding the background label.

### Tissue flow field and strain map analysis

Time-lapse fluorescence image sequences were processed using a custom Python pipeline. Following intensity normalization, cells were segmented independently in each frame using Cellpose (v4.0.7)^62^, and cell centroids were tracked to generate time-resolved cell trajectories. The velocity of each cell was calculated from centroid displacements:

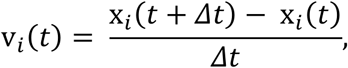

where x_*i*_(*t*) denotes the centroid position of cell *i* at time *t*, and *Δt* is the time interval between frames.

Local tissue deformation was quantified using a cell-centered affine fitting approach. Within a local neighborhood around each target cell, the velocity field was approximated by a first-order affine model:

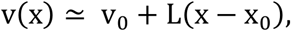

where x_0_ is the position of the target cell, v_0_ is the local translational velocity, and L is the local velocity-gradient tensor:

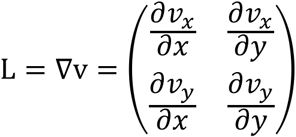

The components of L were estimated by Gaussian-weighted least-squares fitting of neighboring cell velocities, providing a local estimate of spatial velocity variation without binning the velocity field into predefined grids. The strain-rate tensor was obtained as the symmetric part of the velocity gradient:

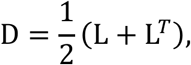

and principal strain rates (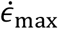 and 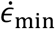) were obtained by eigen decomposition of D; maximum and minimum were defined algebraically, representing the most extensional and most compressive components, respectively. Unless otherwise noted, the term “strain rate” throughout this study refers to the maximum principal strain rate 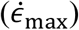.

To quantify accumulated deformation relative to a reference configuration, the local deformation gradient F was estimated using the same weighted fitting framework from cells matched by their tracking identities, and the Green-Lagrange strain tensor:

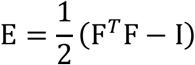

was computed. A scalar equivalent cumulative strain was defined as:

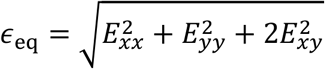

(dimensionless), and the global cumulative strain (relative to frame 0) was obtained by spatially averaging *ϵ*_eq_ over all tracked cells at each time point.

### Correlation analysis

Associations between variables were assessed using the linear mixed-effects model in MATLAB (R2024a, MathWorks). To account for embryo-specific variation and clustering of observations by embryo, embryo identity was included as a random intercept. The general model was specified as:

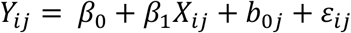

where *X*_*ij*_ and *Y*_*i*j_ denote paired observations *i* from embryo *j, β*_*0*_ is the population-level intercept, *β*_1_ is the fixed-effect regression coefficient, *b*_*0j*_ represents the embryo-specific random intercept, and *ε*_*ij*_ is the residual error. The coefficient *β*_*1*_ quantifies the population-level slope of *Y* on *X* after accounting for embryo-specific variation. Statistical significance was assessed from the fixed-effect term of the mixed-effects model.

### The continuum minimal-response model

The embryo mid-sagittal cross-section is represented by a closed planar curve *r*(*s, t*) = (*∂*(*s, t*), *∂*(*s, t*)) *∈* ℝ^2^, where *s ∈* [0, *L*_0_] is initialized as arc length along the curve at *t* = 0, whose total length is *L*_0_. The local stretch ratio *λ*(*s, t*) and curvature *κ*(*s, t*) in Eq. (1) are defined by

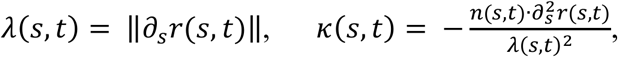

where *n*(*s, t*) is the outward unit normal. The eggshell boundary Γ is parameterized by

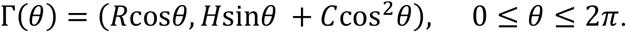

The parameters *R, H, C* are scaled so that 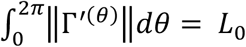.

For the stretching term, we prescribe the target stretch as

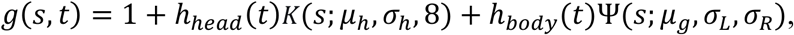

where the head term represents the local strain generated by mitosis, and the body term is an effective target stretch intended to capture the combined influence of GBE-PMG and VF flow on local strain. Their spatial distributions are defined by

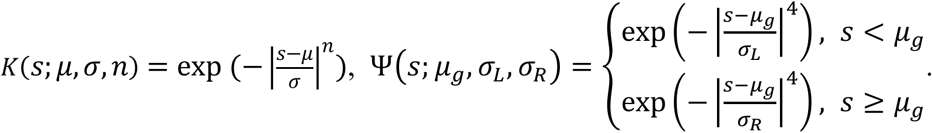

Here, *μ*_*h*_, *μ*_*g*_ specify the region centers, while *σ*_*h*_, *σ*_*L*_, *σ*_*R*_ control their widths. The time-dependent amplitudes follow

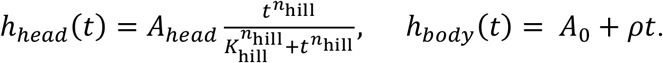

For the bending term, we construct a guide curve *r*_guide_ from a reference curve by prescribing inward displacements at condition-specific dorsal and ventral CF positions on the reference curve *r*_ref_(*s*).

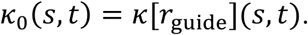

Let *r*_ref_(*S*) denote the arc-length parameterization of the fixed eggshell boundary Γ, with outward unit normal *n*_ref_(*S*). The guide curve is

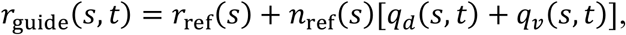

where the inward fold displacements are

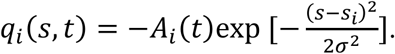

Here, *s*_*d*_, *s*_*v*_ specify the fold positions, *σ* control their width, and *A*_*d*_(*t*), *A*_*v*_ (*t*) prescribe their displacements amplitudes.

We use the same functional forms of *g*(*s, t*) and *κ*_0_(*s, t*) across conditions and represent each perturbation through changes to their prescribed parameters.

The baseline simulation starts at the onset of CF migration (*t* = 0), where *A*_0_ = 0 gives *g*(*s*, 0) = 1 (Fig. 4b) and the two folds are already encoded in *κ*_0_(*s*, 0). Subsequently, the head and body contributions to *g*(*s, t*) increase (Fig. 4c), as do the prescribed fold amplitudes, while the fold centers *s*_*d*_ and *s*_*v*_ used to construct the guided curvature *κ*_0_(*s, t*) remain fixed in material coordinates. Minimizing the energy with these evolving inputs produces dorsal and ventral CF displacements (Fig. 4h, left), measured as shown in Fig. 4d.

For the *stg* RNAi condition, we set *A*_*head*_ = 0, which removes the head contribution to *g*(*s, t*), with all other model parameters unchanged (Fig. 4f). For the 1x *bcd* condition, the dorsal and ventral fold centers *s*_*d*_ and *s*_*v*_ are shifted anteriorly in the construction of *κ*_0_(*s, t*). We also shift and narrow the head component of *g*(*s, t*) through *μ*_*h*_ and *σ*_*h*_, and broaden its body component through *σ*_*R*_ (Fig. 4i). All other parameters retain their baseline values.

### Statistical tests

Statistical analyses were performed using MATLAB (R2024a, MathWorks). Unless otherwise indicated, comparisons between two groups were performed using two-tailed Student’s t-tests. Effect sizes are reported as Cohen’s *d*, calculated using the pooled standard deviation:

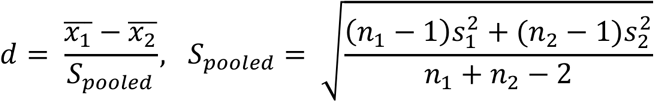

95% confidence intervals (CIs) for Cohen’s *d* were estimated by bootstrap resampling (10,000 iterations, fixed random seed). All statistical tests were two-sided, and labels represent the *P* values: **** < 0.0001, *** < 0.001, ** < 0.01, * < 0.05, and ns > 0.05. Data are presented as mean ± s.d. unless otherwise indicated. The number of biological replicates and the statistical test used for each analysis are specified in the corresponding figure legends.

## Acknowledgements

We thank all members of Dr. Lei-han Tang’s group for the discussions and Dr. Xiaochan Xu for insightful meetings. We thank Dr. Jun Ma and Dr. Feng He for sharing fly stocks and helpful discussions, and Dr. Maoguang Xue and Xiaoyu Liu for their technical support. We are grateful to the Westlake University Microscopy Core Facility for their support, and special thanks to Yinping Gao and Lijie Xu for their support with two-photon microscopy. We thank Dr. Bing He for the suggestion on the manuscript and Dr. Xuan Guo for the fly stocks. Artificial intelligence is used to polish the language and compare methodological options, and the authors remain fully accountable for originality, accuracy, and integrity of the study.

## Funding

This work was supported by the Fundamental and Interdisciplinary Disciplines Breakthrough Plan of the Ministry of Education of China (JYB2025XDXM502) and the National Natural Science Foundation of China (No. 32400668).

## Author contributions

Ruoqing Xu: Conceptualization, Methodology, Software, Validation, Formal analysis, Investigation, Data Curation, Visualization, Writing-original draft, Writing-review and editing; Han Gao: Investigation, Validation; Fancheng Li: Software; Lei-han Tang: Writing-review and editing, Supervision; Hai Zhu: Software, Validation, Writing-review and editing; Zhennan Zhou: Conceptualization, Methodology, Formal analysis, Writing-review and editing; Hanqing Guo: Conceptualization, Methodology, Validation, Data Curation, Writing-original draft, Writing-review and editing, Supervision.

## Code availability

All the models and code generated or used in the study are available in the repository (https://github.com/xrqstart/CF_scripts.git).

## Competing interests

The authors declare no competing interests.

## Figures

**Fig. S1.**
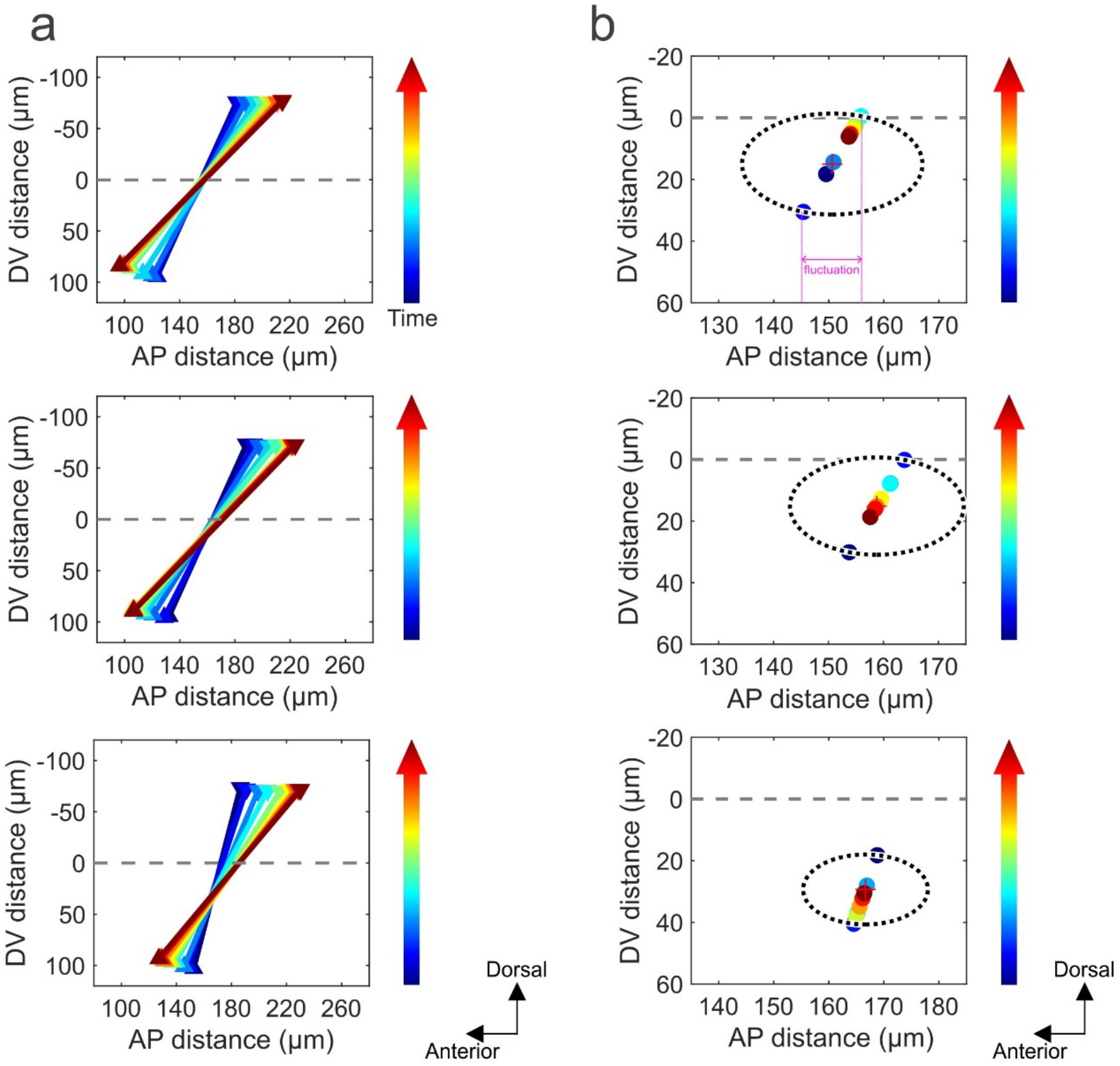
The coherent rotational center of CF migration is maintained over time. **(a)** Absolute positions of the dorsal and ventral CF during the CF migration window. Upward and downward arrows indicate the dorsal and ventral CF positions, respectively. The color gradient from blue to red represents the progression of time. The horizontal dashed line (y = 0) marks the embryo’s AP axis. **(b)** Fluctuation of the instantaneous rotational center, which is defined as the intersection of the CF lines (lines connecting the dorsal and ventral CF) at two consecutive time points. Circled points denote the intersections retained after filtering and used to define the rotational center fluctuations (see Materials and Methods).

**Fig. S2.**
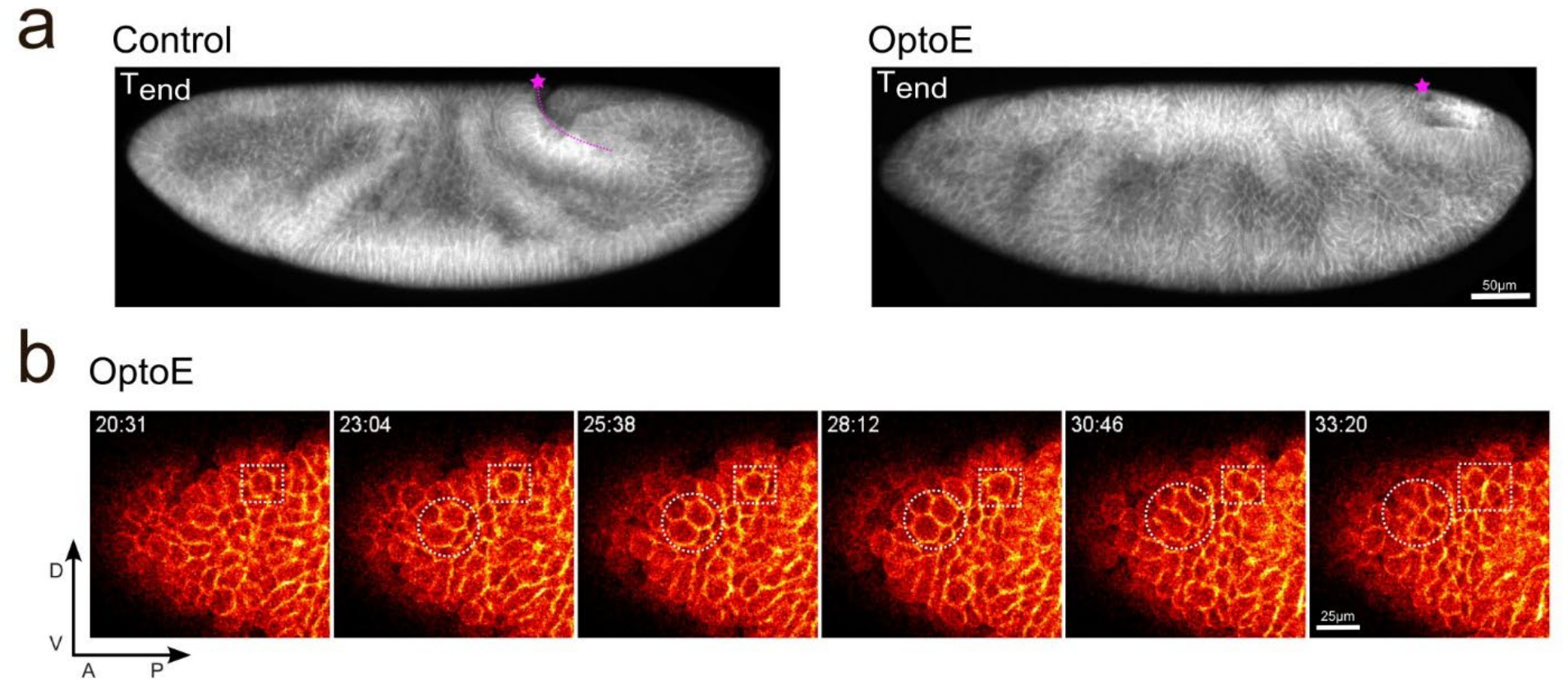
Mitotic activity is preserved in optogenetically inhibited embryos. **(a)** Representative *en face* views of a wild-type and an OptoE embryo at T_end_. The magenta star indicates the position of PMG. The slower movement and abnormal PMG shape in the OptoE embryo indicated the successful inhibition of actomyosin activity. **(b)** Time-lapse images of the head region in the same OptoE embryo. Mitosis is labeled by the dashed circle and box.

**Fig. S3.**
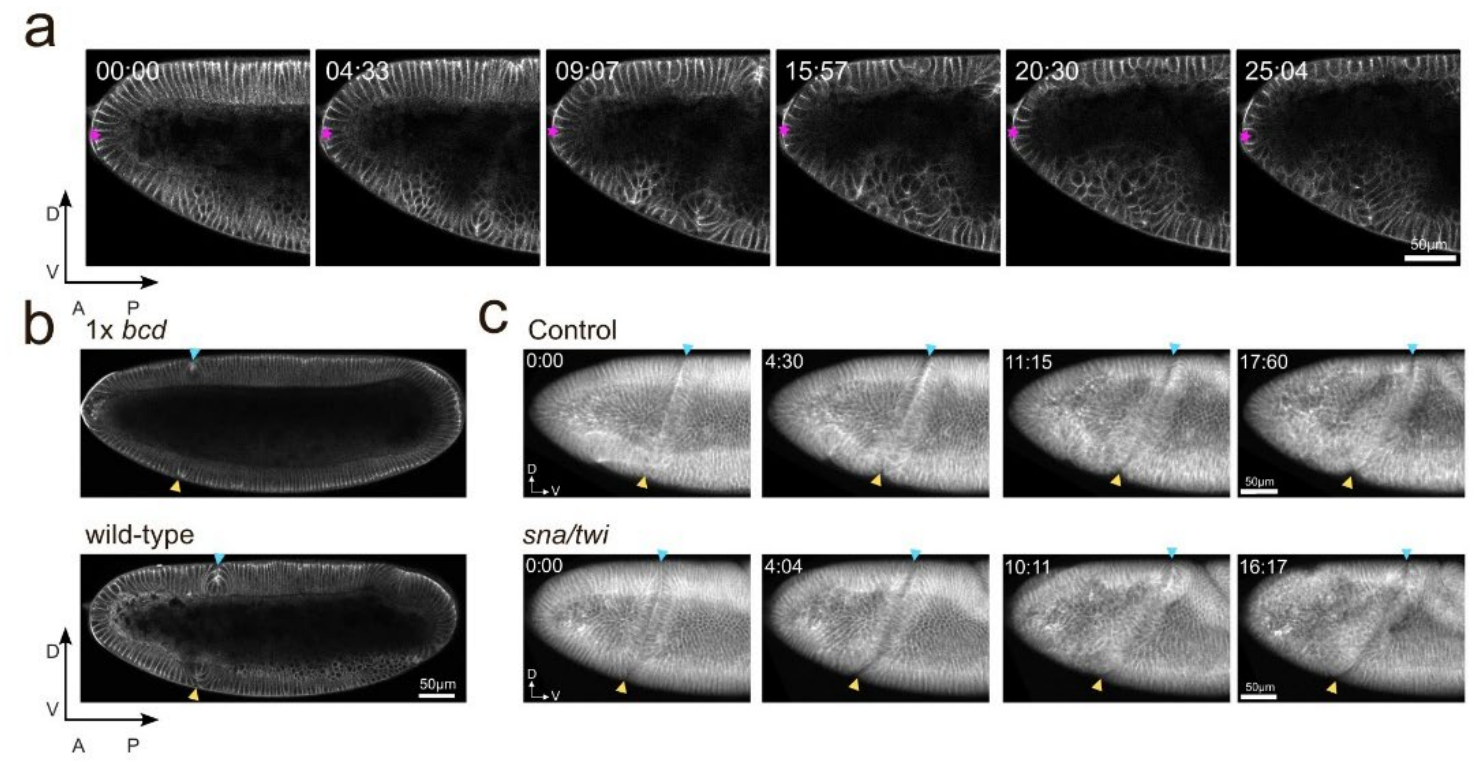
Representative images of embryo morphology and CF dynamics under different conditions. **(a)** The anterior-most cell, marked by the magenta star, was relatively stable during CF migration. **(b)** Representative max mid-sagittal views of maternal 1x *bcd* and wild-type embryos at gastrulation. **(c)** Representative *en face* views of wild-type and *snail/twist* mutant embryos during CF migration. Arrowheads denote CF positions.

## Notes

### Competing Interest Statement

The authors have declared no competing interest.

## References

1. Lee, J.-Y. et al. Wnt/Frizzled signaling controls C. elegans gastrulation by activating actomyosin contractility. Current Biology 16, 1986–1997 (2006).

2. Kölsch, V., Seher, T., Fernandez-Ballester, G. J., Serrano, L. & Leptin, M. Control of Drosophila gastrulation by apical localization of adherens junctions and RhoGEF2. Science 315, 384–386 (2007).

3. Savin, T. et al. On the growth and form of the gut. Nature 476, 57–62 (2011).

4. Bruce, A. E. E. Zebrafish epiboly: Spreading thin over the yolk. Developmental Dynamics 244–258 (2016).

5. Streichan, S. J., Lefebvre, M. F., Noll, N., Wieschaus, E. F. & Shraiman, B. I. Global morphogenetic flow is accurately predicted by the spatial distribution of myosin motors. Elife 7, (2018).

6. Lefebvre, M. F. et al. DynamicAtlas: a morphodynamic atlas for Drosophila development. Nat. Methods 23, 260–270 (2026).

7. Breau, M. A. et al. Extrinsic mechanical forces mediate retrograde axon extension in a developing neuronal circuit. Nat. Commun. 8, (2017).

8. Monnot, P. et al. Intertissue mechanical interactions shape the olfactory circuit in zebrafish. EMBO Rep. 23, e52963 (2022).

9. Hardin, J. & Weliky, M. Cell rearrangement induced by filopodial tension accounts for the late phase of convergent extension in the sea urchin archenteron. Mol. Biol. Cell 30, 1911–1919 (2019).

10. Dzementsei, A., Barooji, Y. F., Ober, E. A. & Oddershede, L. B. Foregut organ progenitors and their niche display distinct viscoelastic properties in vivo during early morphogenesis stages. Commun. Biol. 5, (2022).

11. Nelson, C. M. On buckling morphogenesis. J. Biomech. Eng. 138(2):021005, (2016).

12. Vellutini, B. C. et al. Patterned invagination prevents mechanical instability during gastrulation. Nature 646, 627–636 (2025).

13. Dey, B. et al. Divergent evolutionary strategies pre-empt tissue collision in gastrulation. Nature 646, 637– 646 (2025).

14. Popkova, A. et al. A mechanical wave travels along a genetic guide to drive the formation of an epithelial furrow during Drosophila gastrulation. Dev. Cell 59, 400–414 (2024).

15. Leptin, M. & Grunewald, B. Cell shape changes during gastrulation in Drosophila. Development 110, 73–84 (1990).

16. Martin, A. C., Kaschube, M. & Wieschaus, E. F. Pulsed contractions of an actin–myosin network drive apical constriction. Nature 457, 495–499 (2009).

17. He, B., Doubrovinski, K., Polyakov, O. & Wieschaus, E. Apical constriction drives tissue-scale hydrodynamic flow to mediate cell elongation. Nature 508, 392–396 (2014).

18. Goldner, A. N., Cheikh, M. I., Osterfield, M. & Doubrovinski, K. Viscous shear is a key force in Drosophila ventral furrow morphogenesis. Development 151, (2024).

19. Zallen, J. A. & Wieschaus, E. Patterned gene expression directs bipolar planar polarity in Drosophila. Dev. Cell 6, 343–355 (2004).

20. Bertet, C., Sulak, L. & Lecuit, T. Myosin-dependent junction remodeling controls planar cell intercalation and axis elongation. Nature 429, 667–671 (2004).

21. Blankenship, J. T., Backovic, S. T., Sanny, J. S. S. P., Weitz, O. & Zallen, J. A. Multicellular rosette formation links planar cell polarity to tissue morphogenesis. Dev. Cell 11, 459–470 (2006).

22. Kong, D., Wolf, F. & Großhans, J. Forces directing germ-band extension in Drosophila embryos. Mech. Dev. 144, 11–22 (2017).

23. Costa, M., Wilson, E. T. & Wieschaus, E. A putative cell signal encoded by the folded gastrulation gene coordinates cell shape changes during Drosophila gastrulation. Cell 76, 1075–1089 (1994).

24. Bailles, A. et al. Genetic induction and mechanochemical propagation of a morphogenetic wave. Nature 572, 467–473 (2019).

25. Conte, V. et al. A biomechanical analysis of ventral furrow formation in the Drosophila Melanogaster embryo. PLoS One 7, e34473 (2012).

26. Guo, H., Swan, M. & He, B. Optogenetic inhibition of actomyosin reveals mechanical bistability of the mesoderm epithelium during Drosophila mesoderm invagination. Elife 11, e69082 (2022).

27. Lye, C. M., Blanchard, G. B., Evans, J., Nestor-Bergmann, A. & Sanson, B. Polarised cell intercalation during Drosophila axis extension is robust to an orthogonal pull by the invaginating mesoderm. PLoS Biol. 22, e3002611 (2024).

28. Lye, C. M. et al. Mechanical coupling between endoderm invagination and axis extension in Drosophila. PLoS Biol. 13, e1002292 (2015).

29. Collinet, C., Rauzi, M., Lenne, P. F. & Lecuit, T. Local and tissue-scale forces drive oriented junction growth during tissue extension. Nat. Cell Biol. 17, 1247–1258 (2015).

30. Yu, J. C. & Fernandez-Gonzalez, R. Local mechanical forces promote polarized junctional assembly and axis elongation in Drosophila. Elife 5, e10757 (2016).

31. Driever, W. & Nüsslein-Volhard, C. The bicoid protein determines position in the Drosophila embryo in a concentration-dependent manner. Cell 54, 95–104 (1988).

32. Vincent, A., Blankenship, J. T. & Wieschaus, E. Integration of the head and trunk segmentation systems controls cephalic furrow formation in Drosophila. Development 124, 3747–3754 (1997).

33. Niloy, R. A., Holcomb, M. C., Thomas, J. H. & Blawzdziewicz, J. The mechanics of cephalic furrow formation in the Drosophila embryo. Biophys. J. 122, 3843–3859 (2023).

34. Foe, V. E. Mitotic domains reveal early commitment of cells in Drosophila embryos. Trends in Genetics 5, 322 (1989).

35. Wang, Y. C., Khan, Z. & Wieschaus, E. F. Distinct Rap1 activity states control the extent of epithelial invagination via α-Catenin. Dev. Cell 25, 299–309 (2013).

36. Takeda, M., Sami, M. M. & Wang, Y. C. A homeostatic apical microtubule network shortens cells for epithelial folding via a basal polarity shift. Nat. Cell Biol. 20, 36–45 (2018).

37. Rich, A. & Glotzer, M. Small GTPases modulate intrinsic and extrinsic forces that control epithelial folding in Drosophila embryos. Small GTPases 12, 416–428 (2021).

38. Sabbagh, S., Zhang, H. & Harris, T. J. C. Drosophila anterior midgut internalization via collective epithelial-mesenchymal transition at the embryo surface and enclosure by surrounding tissues. Dev. Biol. 517, 191– 202 (2025).

39. Edgar, B. A. & O’Farrell, P. H. Genetic control of cell division patterns in the Drosophila embryo. Cell 57, 177–187 (1989).

40. Edgar, B. A. & O’Farrell, P. H. The three postblastoderm cell cycles of Drosophila embryogenesis are regulated in G2 by string. Cell 62, 469–480 (1990).

41. Rauzi, M. et al. Embryo-scale tissue mechanics during Drosophila gastrulation movements. Nature Communications 6, 8677 (2015).

42. Prost, J., Jülicher, F. & Joanny, J. F. Active gel physics. Nat. Phys. 11, 111–117 (2015).

43. Pegoraro, A. F., Janmey, P. & Weitz, D. A. Mechanical properties of the cytoskeleton and cells. Cold Spring Harb. Perspect. Biol. 9, a022038 (2017).

44. Leptin, M. twist and snail as positive and negative regulators during Drosophila mesoderm development. Genes Dev. 5, 1568–1576 (1991).

45. Ichbiah, S., Delbary, F., McDougall, A., Dumollard, R. & Turlier, H. Embryo mechanics cartography: inference of 3D force atlases from fluorescence microscopy. Nat. Methods 20, 1989–1999 (2023).

46. Campàs, O. et al. Quantifying cell-generated mechanical forces within living embryonic tissues. Nat. Methods 11, 183–189 (2014).

47. Serwane, F. et al. In vivo quantification of spatially varying mechanical properties in developing tissues. Nat. Methods 14, 181–186 (2017).

48. Garcia, K. E., Kroenke, C. D. & Bayly, P. V. Mechanics of cortical folding: Stress, growth and stability. Philosophical Transactions of the Royal Society B: Biological Sciences 373, 20170321 (2018).

49. Inman, A. et al. A multi-tiered mechanical mechanism shapes the early neural plate. Nature Commun. 16, (2025).

50. Nikolopoulou, E., Galea, G. L., Rolo, A., Greene, N. D. E. & Copp, A. J. Neural tube closure: cellular, molecular and biomechanical mechanisms. Development 144, 552–566 (2017).

51. Kiehart, D. P., Crawford, J. M., Aristotelous, A., Venakides, S. & Edwards, G. S. Cell sheet morphogenesis: Dorsal closure in Drosophila melanogaster as a model system. Annu. Rev. Cell Dev. Biol. 33, 169–202 (2017).

52. Serra, D. et al. Self-organization and symmetry breaking in intestinal organoid development. Nature 569, 66–72 (2019).

53. Fatehullah, A., Tan, S. H. & Barker, N. Organoids as an in vitro model of human development and disease. Nat. Cell Biol. 18, 246–254 (2016).

54. Gjorevski, N. et al. Designer matrices for intestinal stem cell and organoid culture. Nature 539, 560–564 (2016).

55. Brandenberg, N. et al. High-throughput automated organoid culture via stem-cell aggregation in microcavity arrays. Nat. Biomed. Eng. 4, 863–874 (2020).

56. Heisenberg, C. P. & Bellaïche, Y. Forces in tissue morphogenesis and patterning. Cell 153, 948–962 (2013).

57. Wu, X.-L. & Libchaber, A. Particle diffusion in a quasi-two-dimensional bacterial bath. Phys. Rev. Lett. 84, 3017–3020 (2000).

58. Martin, P., Hudspeth, A. J. & Jülicher, F. Comparison of a hair bundle’s spontaneous oscillations with its response to mechanical stimulation reveals the underlying active process. Proc. Natl. Acad. Sci. U. S. A. 98, 14380–14385 (2001).

59. Wilhelm, C. Out-of-equilibrium microrheology inside living cells. Phys. Rev. Lett. 101, 028101 (2008).

60. Mizuno, D., Head, D. A., MacKintosh, F. C. & Schmidt, C. F. Active and passive microrheology in equilibrium and nonequilibrium systems. Macromolecules 41, 7194–7202 (2008).

61. Oda, H. & Tsukita, S. Real-time imaging of cell-cell adherens junctions reveals that Drosophila mesoderm invagination begins with two phases of apical constriction of cells. J. Cell Sci. 114, 493–501 (2000).

62. Pachitariu, M., Rariden, M. & Stringer, C. Cellpose-SAM: superhuman generalization for cellular segmentation. bioRxiv 651001 (2025).

